# 2-Oxoglutarate-dependent dioxygenases contribute to cardenolide biosynthesis in *Erysimum cheiranthoides* (wormseed wallflower)

**DOI:** 10.1101/2025.04.10.648230

**Authors:** Gordon C. Younkin, Cynthia K. Holland, Georg Jander

## Abstract

- Cardenolide biosynthesis evolved convergently in several plant lineages, including wallflowers (*Erysimum*, Brassicaceae). Although the first steps of the biosynthetic pathway, which involve conversion of sterols to pregnane derivatives, have been characterized in *Erysimum cheiranthoides* and other species, key enzymes that catalyze the 14β- and 21-hydroxylation of the steroid core remained elusive.
- In this study, we used comparative transcriptomic analysis of different *Erysimum* species to identify 2-oxoglutarate dependent dioxygenases (2OGDs) that catalyze these reactions in *E. cheiranthoides: Erche03g034150* (*CARDENOLIDE METABOLISM 5*; *CARD5*), which is related to *ALKENYL HYDROXYALKYL PRODUCING 1* (*AOP1*), and *Erche01g020322* (*CARDENOLIDE METABOLISM 6; CARD6*), which arose from a duplication of *DIOXYGENASE FOR AUXIN OXIDATION 1* (*DAO1*).
- Knockout mutants of both genes are deficient in cardenolide biosynthesis and instead accumulate pathway intermediates. Based on transient expression and activity in *Nicotiana benthamiana*, and we identified CARD5 as the likely 14β-hydroxylase, and CARD6 as the 21-oxygenase. Finally, enzyme modeling and substrate docking identified key residues that may allow shifts to substrate recognition during neofunctionalization.
- The results of this research provide new insight into the evolution of cardenolide biosynthesis and have potential practical applications in the engineering of steroid-derived compounds for medical uses.

## Introduction

Cardenolides are a group of steroidal plant specialized metabolites that have fascinated humans for centuries due to their inhibitory activity against animal Na^+^,K^+^-ATPases (Norn & Kruse, 2004). They have evolved convergently in more than 12 plant lineages, including foxglove (*Digitalis*; Plantaginaceae), milkweed (*Asclepias*; Apocynaceae), and wallflowers (*Erysimum*; Brassicaceae)(Agrawal *et al*., 2012). Plant cardenolides primarily function in defense, serving as feeding and oviposition deterrents against insects and other herbivores (Renwick *et al*., 1989; Zalucki *et al*., 2001; Younkin *et al*., 2024a). While these compounds are toxic to humans in high doses, their biological activity makes them useful in the treatment of heart conditions (Norn & Kruse, 2004), and they have been studied more recently for their potential anticancer (Newman *et al*., 2008) and antiviral (Wong *et al*., 2018) properties. Despite their ecological and medical relevance, the full cardenolide biosynthetic pathway has not been described in any plant species.

Cardenolides are derived from steroid metabolism, having a steroid core with an unsaturated 5-member lactone ring on carbon 17, and are often modified via glycosylation, hydroxylation, and acetylation (Agrawal *et al*., 2012). Cardenolide biosynthesis is thought to proceed through pregnane intermediates that are hydroxylated at carbons 14 and 21, with the 14-hydroxy group in the β configuration (Figure 1A) (Kreis, 2017). Importantly, the 14β-hydroxy group, which is characteristic of all cardenolides, is required for their activity against Na^+^,K^+^-ATPases (Schönfeld *et al*., 1985; Bose *et al*., 1988) (Figure 1).

**Figure 1.**
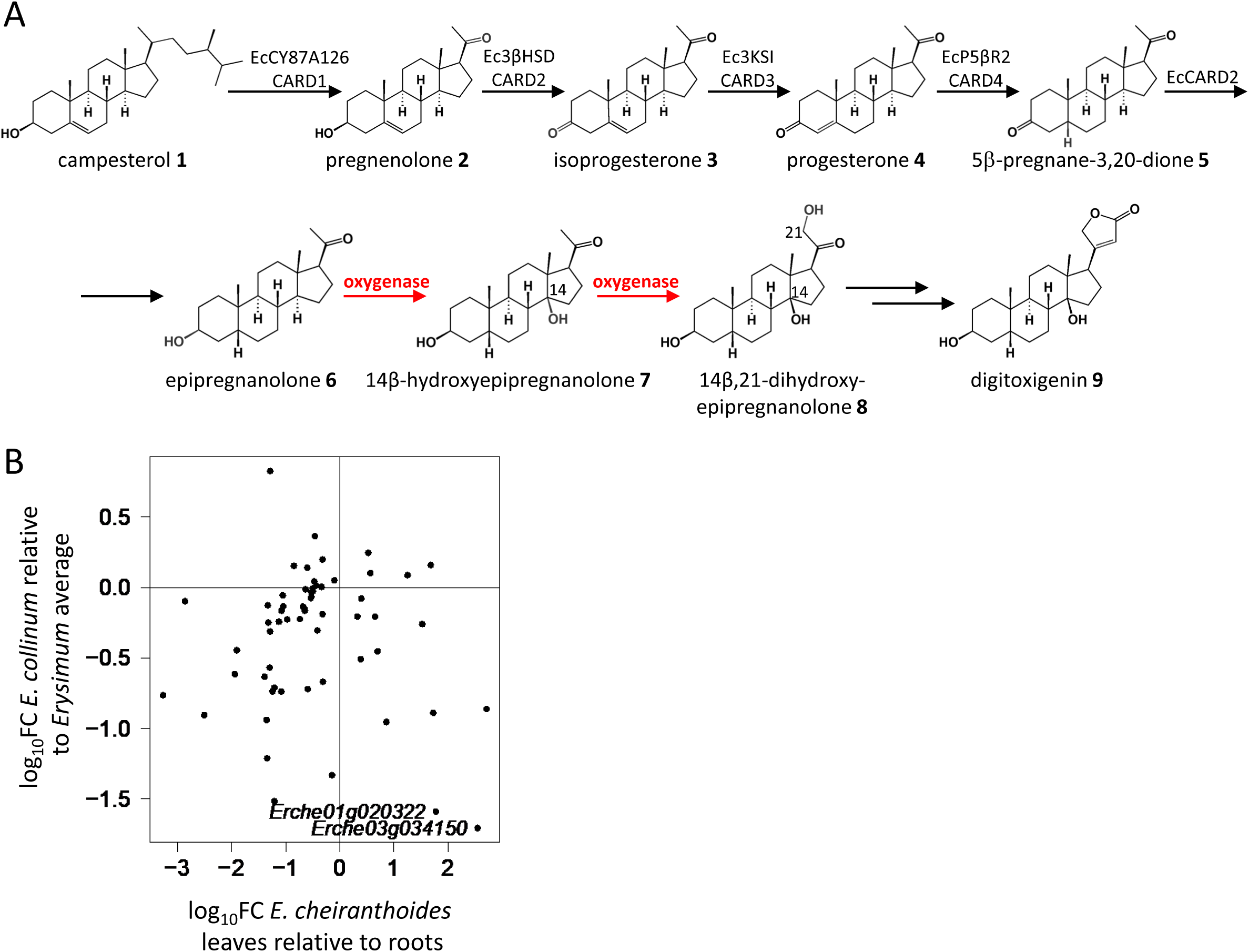
Identification of candidate 2-oxoglutarate dependent dioxygenases (2OGDs) for involvement in cardenolide biosynthesis. (A) Formation and transformation of pregnane intermediates in cardenolide biosynthesis in *Erysimum cheiranthoides*. Previously characterized enzymes are listed, and the steps under investigation in this study, the 14β- and 21-hydroxylation steps, are colored in red. Carbons 14 and 21 are labeled where relevant. (B) Screen of gene expression profiles for *Erysimum* 2OGDs. Genes that are highly expressed in roots relative to leaves and that have low expression in *Erysimum collinum* relative to other *Erysimum* species were of interest. The top candidates selected for further study, *Erche01g020322* and *Erche03g034150*, are indicated. FC: fold-change.

In contrast to earlier steps in the formation of the pregnane intermediates, which have been extensively studied in *Digitalis* and *Erysimum* (Klein *et al*., 2021; Carroll *et al*., 2023; Leykauf *et al*., 2023; Younkin *et al*., 2024b,a; Klein, 2024), no candidate enzymes for the 14β- and 21-hydroxylation have been proposed in any plant species. In animals, pregnane 21-hydroxylation is an essential step in corticosteroid synthesis that is catalyzed by a cytochrome P450, CYP21B (White *et al*., 1986). Mutations to this gene are responsible for 21-hydroxylase deficiency and associated adrenal hyperplasia (Amor *et al*., 1988). However, no analog is known from plants. 14β-hydroxylation of steroids is not frequently reported across the tree of life, with most steroidal compounds being either Δ14-unsaturated or having a 14α substituent group. As such, no enzymes with steroid 14β-hydroxylase activity have been characterized. A cytochrome P450 capable of catalyzing the 14α-hydroxylation of steroids has been identified from the fungal pathogen *Cochliobolus lunatus*, although the regiospecificity of this enzyme is low (Suzuki *et al*., 1993; Chen *et al*., 2019), and the stereochemistry is inverted relative to what is required for cardenolide biosynthesis. The predicted substrate for the 14β-hydroxylase, epipregnanolone **6**, has a 14α-configured hydrogen atom, meaning that stereochemistry must be inverted at that position (Figure 1A). It has been suggested that this unusual inversion of stereochemistry at carbon 14 may involve a multi-step process proceeding through Δ8,14-unsaturated intermediates. However, feeding of such intermediates in *Digitalis purpurea* supports a pathway where 14β-hydroxylation and stereochemical inversion occur in a single step (Deluca *et al*., 1987).

Although there are no characterized plant enzymes with steroid 14β- and 21-hydroxylase activity, plant cytochromes P450 (P450s) and 2-oxoglutarate-dependent dioxygenases (2OGDs) are known to catalyze the oxidation of steroids and triterpenoids at other positions. P450s, which constitute the largest family of metabolic enzymes in plants and represent approximately 1% of plant protein-coding genes, catalyze diverse oxidation reactions in both primary and specialized metabolism (Nelson & Werck-Reichhart, 2011). They have long been known to act on varied triterpenoid substrates including α- and β-amyrin (Yasumoto *et al*., 2016), cucurbitacins (Zhou *et al*., 2016), and brassinosteroids (Ohnishi *et al*., 2009). Although several P450s involved in cardenolide biosynthesis have been discovered in *Digitalis lanata* (Carroll *et al*., 2023), *Calotropis procera* (Kunert *et al*., 2023), and *Erysimum cheiranthoides* (Younkin *et al*., 2024a), none of these enzymes catalyze the 14β- or 21-hydroxylation of cardenolide intermediates.

2OGDs are the second-largest enzyme family in plants and, like P450s, contribute to a wide variety of metabolic pathways (Hagel & Facchini, 2018). However, the importance of 2OGDs in the oxidation of steroidal metabolites was discovered only recently, with a 16α-hydroxylase (Nakayasu *et al*., 2017; Hagel & Facchini, 2018) and a 23-hydroxylase (Nakayasu *et al*., 2020) involved in steroidal glycoalkaloid metabolism in *Solanum*. Here, we screened *E. cheiranthoides* 2OGDs to identify predicted 14β- and 21-hydroxylases that are required for cardenolide biosynthesis. *Erche03g034150* is designated as *CARDENOLIDE METABOLISM 5* (*CARD5*), and *Erche01g023022* is designated as *CARDENOLIDE METABOLISM 6* (*CARD6*). CRISPR/Cas9 knockouts of both enzymes are deficient in cardenolide biosynthesis, and a combination of substrate feeding, transient expression, and enzyme modeling support their function as steroid 14β- and 21-oxygenases.

## Materials and Methods

### Plants and growth conditions

A genome-sequenced isolate of *E. cheiranthoides* (Züst *et al*., 2020), Arabidopsis Biological Resource Center (https://abrc.osu.edu) accession number CS29250, was used for all experiments. Plants were grown in Cornell Mix (by weight 56% peat moss, 35% vermiculite, 4% lime, 4% Osmocote slow release fertilizer [Scotts, Marysville, OH], and 1% Unimix [Scotts]) in Conviron (Winnipeg, CA) growth chambers with a constant temperature of 23 °C, and 180 µM m ^-2^ s^-1^ photosynthetic photon flux density, and a 16:8 h day:night cycle.

### RNA-sequencing, cloning, and knockout of candidate genes

Raw RNA-sequencing reads from 48 *Erysimum* species (Züst *et al*., 2020) (PRJNA563696) and various *E. cheiranthoides* tissues (Younkin *et al*., 2024a) (PRJNA1015726) were downloaded from the NCBI Short Read Archive. Read counts were quantified and analyzed as described previously (Younkin *et al*., 2024a), except that the *E. cheiranthoides* genome annotation v2.1 was used (Mirzaei *et al*., 2024). Cloning of candidate genes, and CRISPR/Cas9 knockout in *E. cheiranthoides* were performed as described previously (Younkin *et al*., 2024a). Primers used for cloning candidate genes, generating CRISPR/Cas9 mutants, and screening mutations are provided in Table S1.

### Substrate feeding and transient expression

Genes were transiently expressed in leaves of 4-week-old *Nicotiana benthamiana* plants. *Agrobacterium tumefaciens* growth and infiltration were as described previously (Younkin *et al*., 2024a). For pathway reconstruction, previously characterized cardenolide biosynthetic enzymes *CARD1*, *CARD2*, *CARD3*, and *CARD4* were expressed together and with either *CARD5* only or both *CARD5* and *CARD6*. For substrate feeding experiments, *CARD5* and *CARD6* were infiltrated individually and as a pair, and separate plants expressing GFP were used as a negative control. A 200 μM solution of pregnenolone (Sigma-Aldrich, St. Louis, MO), progesterone (Sigma-Aldrich), or epipregnanolone (Cayman Chemical, Ann Arbor, MI, USA) in 10 mM 2-(N-morpholino)ethanesulfonic acid (MES) buffer and 10 mM MgCl_2_ were infiltrated into leaves expressing candidate genes three days after *A. tumefaciens* infiltration, with three replicates for each gene/substrate combination and buffer-only controls. For *CARD6*, a 50% methanol extract of *card6* knockout lines, diluted 10x into MES buffer, was infiltrated to test whether the intermediates accumulating in the mutant could serve as substrates. Tissue was collected four days after *A. tumefaciens* infiltration for LC-MS analysis.

We supplemented *E. cheiranthoides* mutant lines with intermediates in cardenolide biosynthesis to test for restoration of the pathway. A 200 μM solution of progesterone or 21-hydroxyprogesterone (Sigma-Aldrich) in MES buffer (see above), was infiltrated into the abaxial surface of young leaves of three-week old plants of *card5-1* and *card6* mutant lines, with three replicates per substrate/mutant combination. Tissue was collected after 24 hours for LC-MS analysis.

### Liquid chromatography-mass spectrometry (LC-MS) analysis

Methanolic extracts of *E. cheiranthoides* and *N. benthamiana* leaves (Younkin *et al*., 2024a) were analyzed on an UltiMate 3000 UHPLC system coupled to a Q-Exactive hybrid quadrupole-orbitrap mass spectrometer (ThermoFisher Scientific). The instrument was fitted with a Supelco Titan^TM^ C18 UHPLC Column (80Å, 100 x 2.1 mm, particle size 1.9 μm; Sigma Aldrich). Injections of 2 μL were separated by a solvent gradient consisting of mobile phase A (water + 0.1% (v/v) formic acid) and mobile phase B (acetonitrile + 0.1% (v/v) formic acid): 0-0.5 minutes, hold at 2% B; 0.5-10 minutes, linear gradient from 2%-97% B; 10-11.5 minutes, hold at 97% B, 11.5-13 minutes, hold at 2% B. All solvents were Optima LC/MS grade (ThermoFisher Scientific). The solvent flow rate was 0.5 mL/minute, the column oven was set to 40 °C, and the autosampler temperature was 15 °C. The mass spectrometer was run in full scan positive ionization mode. Targeted MSMS spectra were collected with an isolation window of 2.0 *m/z* and normalized collision energy of 30%.

Untargeted analysis of LC-MS data was conducted using XCMS (Smith *et al*., 2006; Tautenhahn *et al*., 2008; Benton *et al*., 2010) in R statistical software (R Core Team, 2020) with the following functions and parameters. CentWave: snthresh 3, peakwidth c(2,7), ppm 2.5, noise 10000, prefilter c(3, 10000); MergeNeighboringPeaks: expandRt 2; Obiwarp: binSize 0.01; PeakDensity: minFraction 0.5, bw 2, binSize 0.01, and FillChromPeaks with default parameters. Once chromatographic peaks of interest were identified, peak areas were re-quantified in a targeted manner using a custom processing method in Xcalibur^TM^ Software (ThermoFisher Scientific) with the following parameters: peak detection ICIS, smoothing points 1, baseline window 40, area noise factor 5, peak noise factor 15, tailing factor 2. Mass features used for quantification are provided in Table S2 for cardenolides and compounds **10**-**11** found in *E. cheiranthoides* mutants, and Table S3 for compounds **12**-**21** from *N. benthamiana* assays. Authentic standards were not available for the hypothesized structures of compounds **12**-**21**, but a 70% methanolic extract of *Cynanchum paniculatum* (Paniculate Swallowwort Root; KHT Herbs & Goods, Sacramento, CA) was used to validate identification of compound **12** as 14β-hydroxypregnenolone.

### Statistical and phylogenetic analysis

For statistical comparisons between compound abundances in mutant lines and transient expression assays, the aov and TukeyHSD functions in R statistical software (R Core Team, 2020) were used for one-way ANOVA and post-hoc Tukey’s HSD tests. Log-transformed peak areas that had been normalized to the internal standard were used for analysis. Plots were made using MSnbase (Gatto & Lilley, 2012; Gatto *et al*., 2021), and multcompView (Graves *et al*., 2023).

Sequences homologous to *CARD5* and *CARD6* were identified using BLAST against publicly available transcriptomes for *Arabidopsis thaliana* (Berardini *et al*., 2015) and *E. cheiranthoides* (Züst *et al*., 2020) (NCBI PRJNA563696). For *CARD5*, a functionally characterized 2OGD from *Solanum lycopersicum*, *Sl23DOX*, was also included (Nakayasu *et al*., 2020). In the case of *CARD6*, orthologs were retrieved from *Oryza sativa* (Kawahara *et al*., 2013), *Brassica oleracea*, *Camelina sativa*, and *Eutrema salsugineum* (Tello-Ruiz *et al*., 2022). Outgroups were selected from sister 2OGD clades based on a comprehensive phylogenetic analysis of plant 2OGDs (Kawai *et al*., 2014). Nucleotide sequences were aligned using ClustalW (Sievers *et al*., 2011; Madeira *et al*., 2022). Gene phylogenies were inferred using the IQ-TREE web server (Trifinopoulos *et al*., 2016; Hoang *et al*., 2018; Minh *et al*., 2020) with default parameters, except that bootstrap alignments were increased to 10,000. Raw data underlying all figures are available in the Supporting Information.

### Enzyme modeling and substrate docking

Three-dimensional structural models of each enzyme were obtained from AlphaFold (Jumper *et al*., 2021) (AtDAO1: Q9XI75) or were generated using the amino acid sequence of the protein using ColabFold v1.5.5 (Mirdita *et al*., 2022). Metal ions and alpha-ketoglutarate were added to each protein using AlphaFill (Hekkelman *et al*., 2023). Because AtDAO1 was co-crystalized with Mg ions, these are included in the DAO1 models. Substrate structures (indole-3-acetic acid, a diastereomer of **11**, glucoiberin, and pregnenalone) were obtained from Zinc20 (zinc.docking.org), and were docked into enzyme active sites using AutoDock Vina (ver. 1.1.2) with grid box dimensions of 40 x 40 x 40 Å and an exhaustiveness of 8 (Trott & Olson, 2010; Forli *et al*., 2016). Docking results were visualized in PyMOL (ver. 2.5.7) (https://www.pymol.org/2/).

## Results

### Identification of candidate 2OGDs

We examined the expression patterns of *Erysimum* 2OGDs to identify candidates for involvement in cardenolide biosynthesis. Because cardenolides are synthesized in *E. cheiranthoides* leaves and transported to the roots (Alani *et al*., 2021), we expect cardenolide biosynthetic genes to be highly expressed in leaves compared to roots. Furthermore, cardenolide biosynthesis has been lost in *Erysimum collinum* (Züst *et al*., 2020), so we expect low expression in *E. collinum* compared to all other species of *Erysimum*. Using an approach that we previously applied for screening P450s (Younkin *et al*., 2024a), we searched for 2OGDs following these expression patterns and identified two genes, *Erche01g020322* and *Erche03g034150*, as top candidates (Figure 1B). Notably, *Erche03g034150* clustered with other cardenolide biosynthesis genes in a coexpression analysis across 48 *Erysimum* species (Younkin *et al*., 2024b).

### Phylogenetic analysis of candidate genes

To better understand the phylogenetic context of the candidate 2OGDs and the activity of related enzymes, we inferred phylogenetic trees for each gene and examined microsynteny between the genomic regions of *E. cheiranthoides* and *A. thaliana* containing these genes. *Erche01g020322* is a tandem duplicate of *DIOXYGENASE FOR AUXIN OXIDATION 1* (*EcDAO1*) (Figure 2A-B), placing it within 2OGD clade DOXC15 (Kawai *et al*., 2014). *AtDAO1* and the closely related *AtDAO2* have a critical role in auxin homeostasis through its oxidative deactivation (Zhao *et al*., 2013; Porco *et al*., 2016; Zhang *et al*., 2016). In *E. cheiranthoides*, *EcDAO1* has been duplicated relative to *A. thaliana* (Figure 2A,B). This duplicate copy, *Erche01g020322*, sits on a relatively long branch in the DAO phylogenetic tree, suggestive of rapid evolution of the gene sequence, with potential for neofunctionalization of the encoded protein.

**Figure 2.**
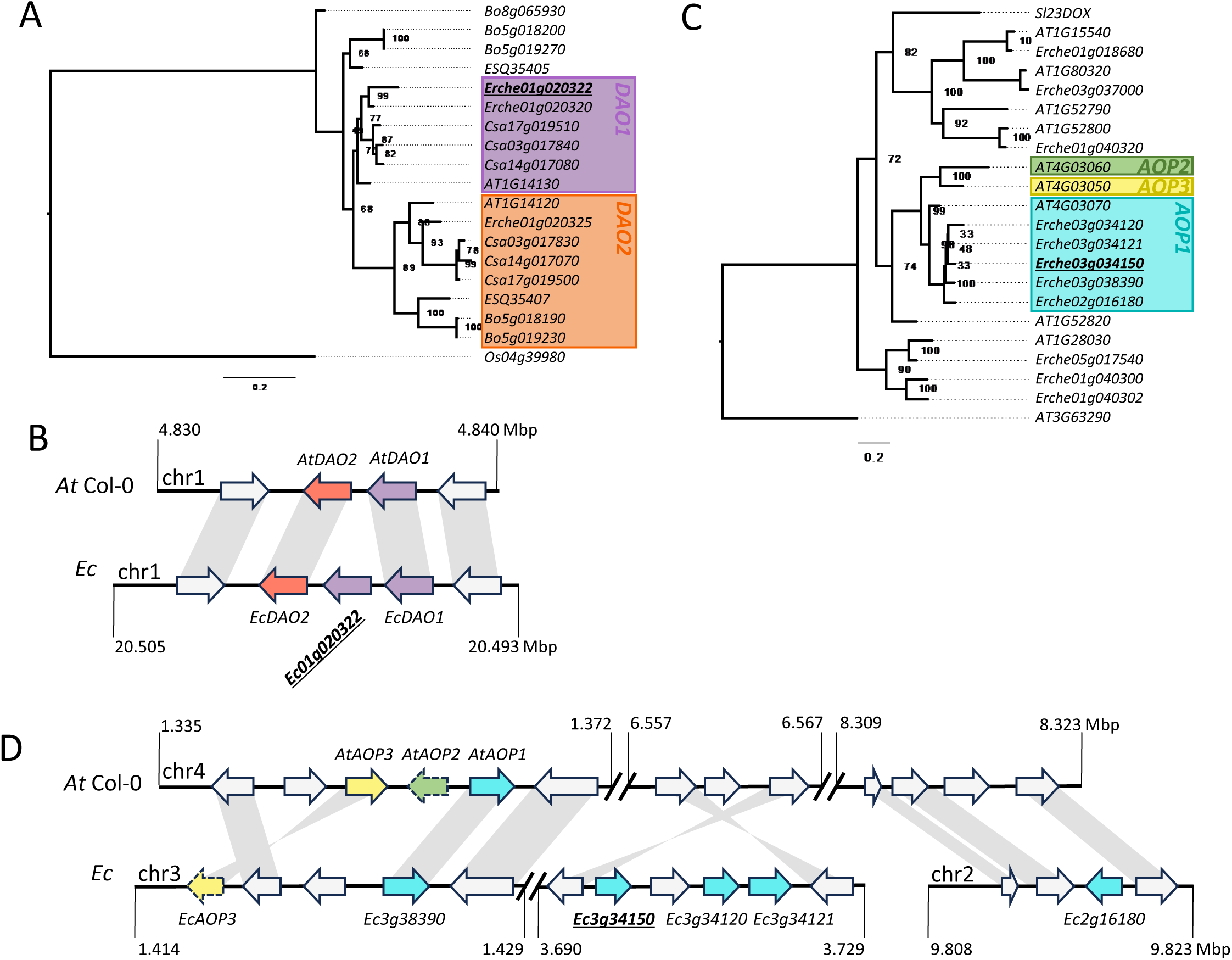
Phylogenetic analysis of candidate genes. Nucleotide phylogenies were inferred from coding sequences of 2-oxoglutarate-dependent dioxygenase (2OGD) clades (A) DOXC15 and (C) DOXC20 from *Arabidopsis thaliana* Col-0 (At) and *Erysimum cheiranthoides* (Ec). Candidate genes for involvement in cardenolide biosynthesis are underlined and in bold. Select genes from other species: *Brassica oleracea* (Bo), *Camelina sativa* (Csa), *Eutrema salsugineum* (ES), *Oryza sativa* (Os), and *Solanum lycopersicum* (Sl). Scale bars indicate estimated number of substitutions per site, numbers at nodes are bootstrap support values. (B) and (D): Microsynteny plots comparing loci containing candidate 2OGDs in *E. cheiranthoides* (Ec) with syntenic regions in *A. thaliana* Col-0 (At). Pseudogenes are indicated with a dashed border. Colors from gene phylogenies are carried over into the microsynteny plots.

*Erche03g034150* belongs to the DOXC20 group of plant 2OGDs (Kawai *et al*., 2014), and is nested within the AOP (akenyl hydroxyalkyl producing) clade. In *A. thaliana*, AtAOP2 and AtAOP3 have been implicated in the modification of aliphatic glucosinolates (Kliebenstein *et al*., 2001), with structural variation of the locus explaining variation in glucosinolate profiles among *A. thaliana* accessions (Chan *et al*., 2010). *AtAOP1*, which is the phylogenetically most closely related *A. thaliana* gene to *Erche03g034150* (Figure 2C), does not have a known function. Compared to the single copy of *AOP1* in *A. thaliana*, which is contained within the GS-AOP locus (Kliebenstein *et al*., 2001), there are five copies in *E. cheiranthoides* that are scattered across three clusters on chromosomes 2 and 3 (Figure 2D). It should be noted that *A. thaliana* chromosome 4 is syntenic with parts of *E. cheiranthoides* chromosomes 2 and 3 (Züst *et al*., 2020), so this broad genomic distribution may be explained by tandem duplication followed by chromosomal fission.

### Knockout of candidate genes

We generated CRISPR/Cas9 knockouts of both candidates to assess their potential roles in cardenolide biosynthesis. Two independent knockout mutants were produced for *Erche03g034150* (*card5-1* and *card5-2*), while only one knockout line was recovered for *Erche01g020322* (*card6*). Sequences of the mutant alleles are available in Figures S1-S2. Knockout lines for both genes were deficient in cardenolide biosynthesis, with a ∼1,000-fold reduction in total cardenolide-related peak area for both *card5* lines, and a ∼100-fold reduction for *card6* (Figure 3A, Table S4-S5). In knockout mutants, the accumulation of pathway intermediates can provide clues as to the role a gene plays in a pathway. In *card5-1* and *card5-2*, pregnenolone and progesterone accumulated to higher levels than in wildtype or *card6* plants (Figure 3B,C), suggesting that CARD5 may be the first hydroxylase to act on these pregnane intermediates. In *card6*, two apparently hydroxylated pregnane intermediates accumulate to very high levels (Figure 3D-F, Table S4-S5). The masses and fragmentation patterns of these compounds are consistent with either mono- (compound **11**), or di- (compound **10**) glycosylated hydroxyepipregnanolone (Figure 3G,H, S3). Notably, though compound **11** occurs at trace levels in wildtype plants, it is even less abundant in *card5-1* and *card5-2* plants (Figure 3E, Table S4-S5).

**Figure 3.**
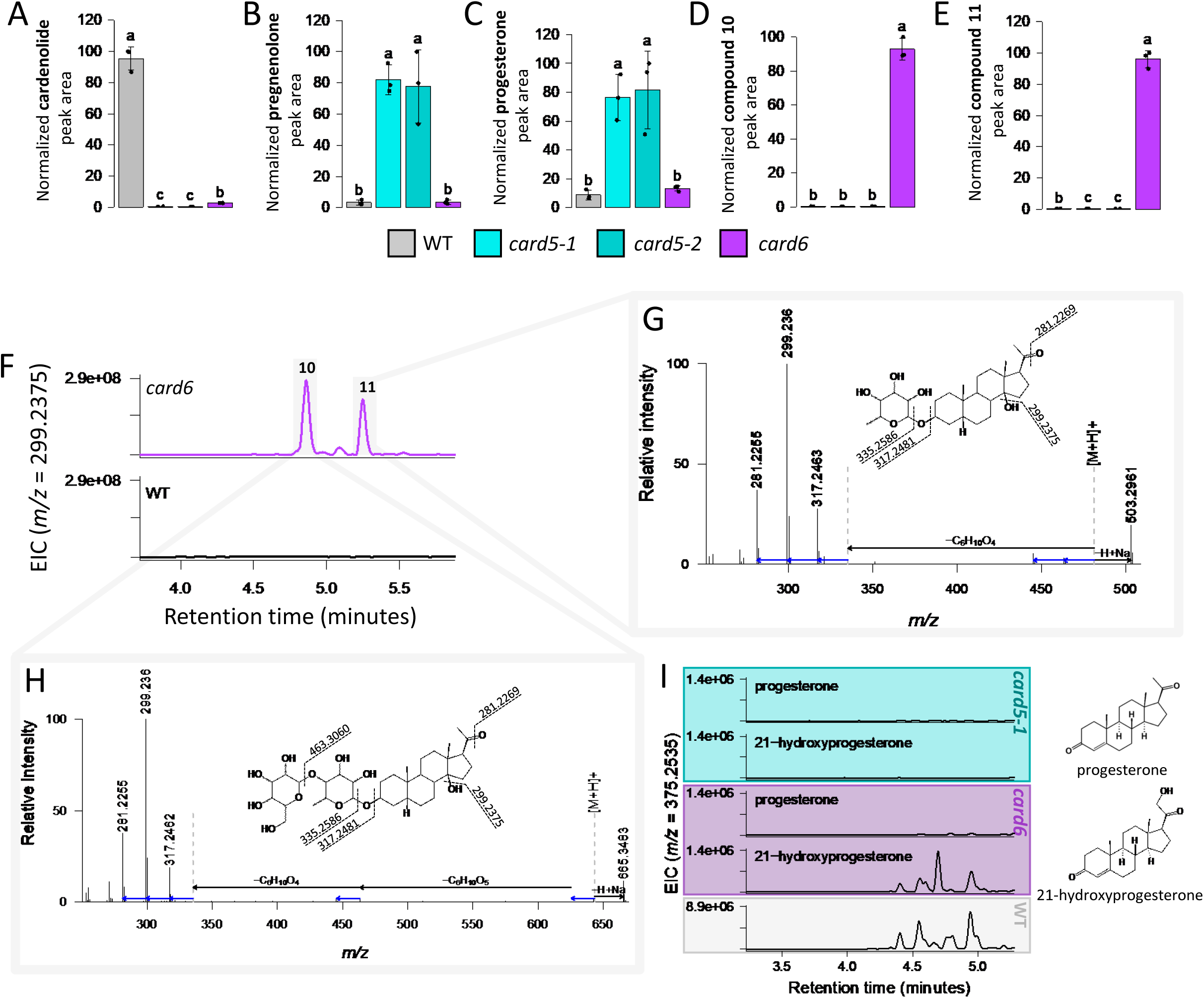
Characterization of mutant lines. Normalized peak area for (A) total cardenolides, (B) pregnenolone, (C) progesterone, (D) compound **10**, and (E) compound **11** from wildtype *Erysimum cheiranthoides* (WT) and CRISPR/Cas9 mutants of *Erche03g034150* (*card5-1* and *card5-2*, cyan) and *Erche01g020322* (*card6*, magenta). N=3 replicates per line. Letters indicate differences between groups, *P*<0.05, one-way ANOVA with post-hoc Tukey’s HSD on log-transformed data. Full results of statistical tests are available in Table S5. Error bars are ± s.d. (F) Extracted ion chromatogram (EIC) of *m/z*=299.2375, a fragment common to compounds **10** and **11**. Full MS scans for (G) compound **11** and (H) compound **10**, with hypothetical structure based on pattern of neutral losses. Hypothetical [M+H]^+^ and [M-sugar]^+^ peaks, which were not detected, are shown as grey dashed lines. (I) EIC of *m/z*=375.2535, a fragment common to all digitoxigenin glycosides, following infiltration of leaves of mutant plants with cardenolide intermediates. Chromatograms are representative samples from from N=3 replicates.

These results are consistent with a pathway where CARD5 acts first to hydroxylate epipregnanolone **6**, followed by a second hydroxylation by CARD6. However, the knockout lines alone do not confirm the regiospecificity of these enzymes. In order to address this question, we fed mutant lines with both progesterone and 21-hydroxyprogesterone. Progesterone did not restore cardenolide biosynthesis in either line, but 21-hydroxyprogesterone did restore cardenolide biosynthesis in *card6* (Figure 3I). This is strong evidence that CARD6 is required for steroid 21-hydroxylation. While we lack direct evidence for the specific activity of CARD5, by process of elimination it is most likely the steroid 14β-hydroxylase.

### Transient expression in Nicotiana benthamiana

In order to further investigate the activity of CARD5 and CARD6, we transiently expressed the genes encoding these enzymes in *N. benthamiana* leaves, co-infiltrated potential substrates, and monitored activity via LC-MS. CARD5 hydroxylated all substrates we provided it with, including pregnenolone, progesterone, and epipregnanolone, to form compounds **12**, **13**, and **14** (Figures 4A-C, S4, Table S6-S7). Based on the results of substrate feeding to *E. cheiranthoides* mutant lines, we predict that these products are 14β-hydroxylated. While authentic standards were not available for these compounds, **12** is found in *Cynanchum paniculatum* (Sugama *et al*., 1986), which we used to confirm the identification of **12** as 14β-hydroxypregnenolone (Figure S4). Consistent with evidence from the mutant lines that CARD5 catalyzes the first hydroxylation step, we did not detect CARD6 activity directly on any of the substrates provided (Figure 4A-C, Table S6-S10). However, when we expressed CARD6 together with CARD5, we observed consumption of compounds **12**, **13**, and **14** and formation of new compounds of unknown structure, which we designate as compounds **15**-**20** (Figure 4D-F, Table S6-S7). Based on similar *m/z*, predicted molecular formula, and MSMS fragmentation, these compounds appear to be related, with compounds **15** and **16** likely retaining the Δ5-double bond from pregnenolone, compounds **17** and **18** retaining the α,β-unsaturated ketone from progesterone, and compounds **19** and **20** having a fully saturated steroid core (Figures 4D-F, S5). As expected, these compounds contain an extra oxygen atom relative to compounds **11**-**13**, but their *m/z* is 2 Da less than would be expected for hydroxylated derivatives, suggesting that the hydroxy group may have been further oxidized to a carbonyl. Furthermore, these compounds appear to be attached to a large substituent group with a predicted molecular formula C_9_H_12_O_8_, which might correspond to a malonyl-hexose group. These substituents are presumably added by endogenous *N. benthamiana* enzymes. Hypothetical structures for these compounds are provided in Figure 4D-F. For both CARD5 and CARD6, the compounds identified here were some of the most abundant products, but they were not the only products. The accumulation of other compounds with similar masses and retention times, perhaps representing further hydroxylation or glycosylation of the intermediates described here, was observed in all cases (Tables S8-S10).

**Figure 4.**
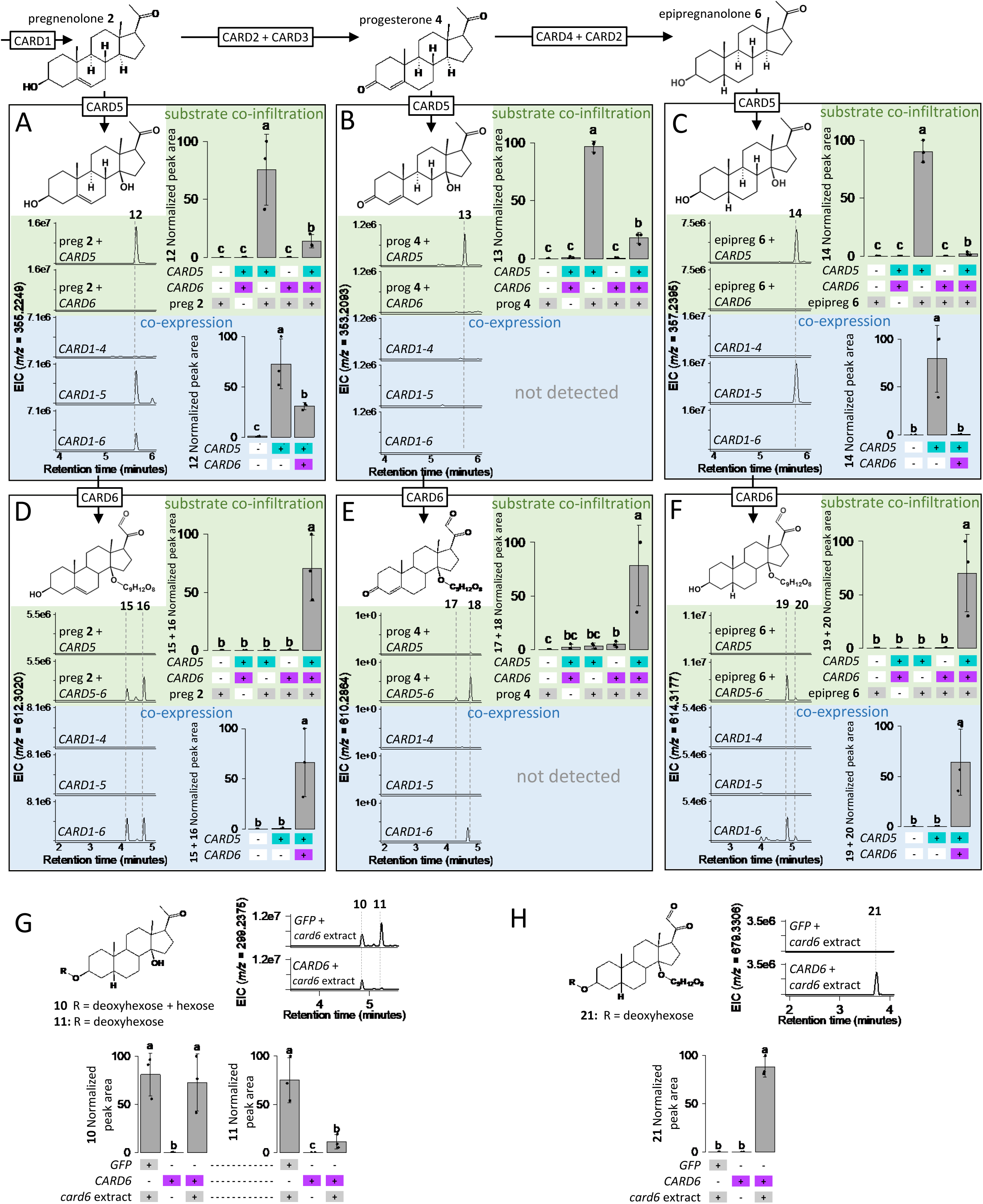
Co-infiltration of *CARD5* and *CARD6* with cardenolide intermediates in *Nicotiana benthamiana*. (A-H) Extracted ion chromatograms (EIC) from representative samples and bar plots showing compounds detected following substrate co-infiltration (green background) or co-expression with upstream pathway enzymes *CARD1, CARD2, CARD3,* and *CARD4* (blue background). All chromatograms show [M+Na]^+^ for each compound, as the sodium adducts were the most reliably detected ions. For G-H, a diluted methanolic extract from *card6* mutants was co-infiltrated with *CARD6*. Bar plots show peak areas from N=3 replicates per enzyme/substrate combination. Letters indicate differences between groups, *P*<0.05, one-way ANOVA with post-hoc Tukey’s HSD on log-transformed data. Full results of statistical test are available in Tables S7 and S11. Error bars are ± s.d. Abbreviations: pregnenolone (preg), progesterone (prog), epipregnanolone (epipreg). Predicted molecular structures are displayed alongside chromatograms. Structures for compounds **13-21** are hypotheses that have not been confirmed by comparison to authentic standards. Compound **12** was validated by comparison with *Cynanchum paniculatum* extracts (Figure S4).

Next, we infiltrated a methanolic extract from the *card6* mutant line together with *CARD6* into *N. benthamiana* to test whether the enzyme was active on the glycosylated intermediates found in *card6* plants (Figure 2F). Interestingly, we observed consumption of mono-glycosylated compound **11**, but not of the di-glycosylated compound **10** (Figure 4G, Tables S6-S7), and saw the accumulation of compound **21**, which was similar to compounds **15**-**20** (Figures 4H, S5, Tables S6-S7). Finally, we expressed CARD5 and CARD6 with the first four characterized genes from the *E. cheiranthoides* cardenolide biosynthesis pathway, which we designate *CARDENOLIDE METABOLISM 1* – *CARDENOLIDE METABOLISM 4*: *EcCYP87A126* (*CARD1*), *Ec3βHSD* (*CARD2*), *Ec3KSI* (*CARD3*), and *EcP5βR2* (*CARD4*) in an attempt to reassemble part of the pathway in a heterologous system. When only *CARD1*-*CARD5* were present, we observed accumulation of compounds **12** and **14**, and when *CARD6* was included, compounds **15**, **16**, **19**, and **20** were formed (Figures 4A-F, Tables S6, S11).

### Enzyme modeling and substrate docking

To understand the evolution of CARD5 and CARD6 in *Erysimum*, we generated structural models of each protein to visualize their active sites with their predicted substrates. When the global architecture of the monomers was compared, we noted a large stretch of 95 amino acids between residues 176 and 271 that are disordered in the AtAOP3 model and entirely absent in the CARD5 protein (Figure 5A). In each of their active sites, the Fe(II) ion is coordinated with the canonical His/Asp/His triad and 2-oxoglutarate (2OG) is bound (Figure 5B, C). Glucoiberin (3-(methylsulfinyl)propylglucosinolate) was docked into the AOP3 active site, and pregnenalone was docked into the active site of CARD5, which allowed us to compare the putative active sites of these two enzymes. While a number of active site residues are conserved between the two enzymes (Pro 82, Phe 83, Leu 89, Ser 94, Arg 166, Met 168, Leu 183, His 186, Thr 187, Asp 188, Lys 189, His 243, Ala 257, Phe 259, Phe 288, His 296, and Arg 301 in CARD5), there are ten residues that vary. Two residues that have nonpolar side chains in AtAOP3 (Leu 85 and Phe 386) are also nonpolar in CARD5 (Phe 87 and Met 292, respectively). Notably, three polar residues in the AtAOP3 active site (Arg 162, Asn 355, and Tyr 389) are nonpolar residues in CARD5 (Met 164, Leu 261, and Phe 295, respectively). This presence of nonpolar active site residues in CARD5 active site may account for the activity with steroid substrates like pregnenalone.

**Figure 5.**
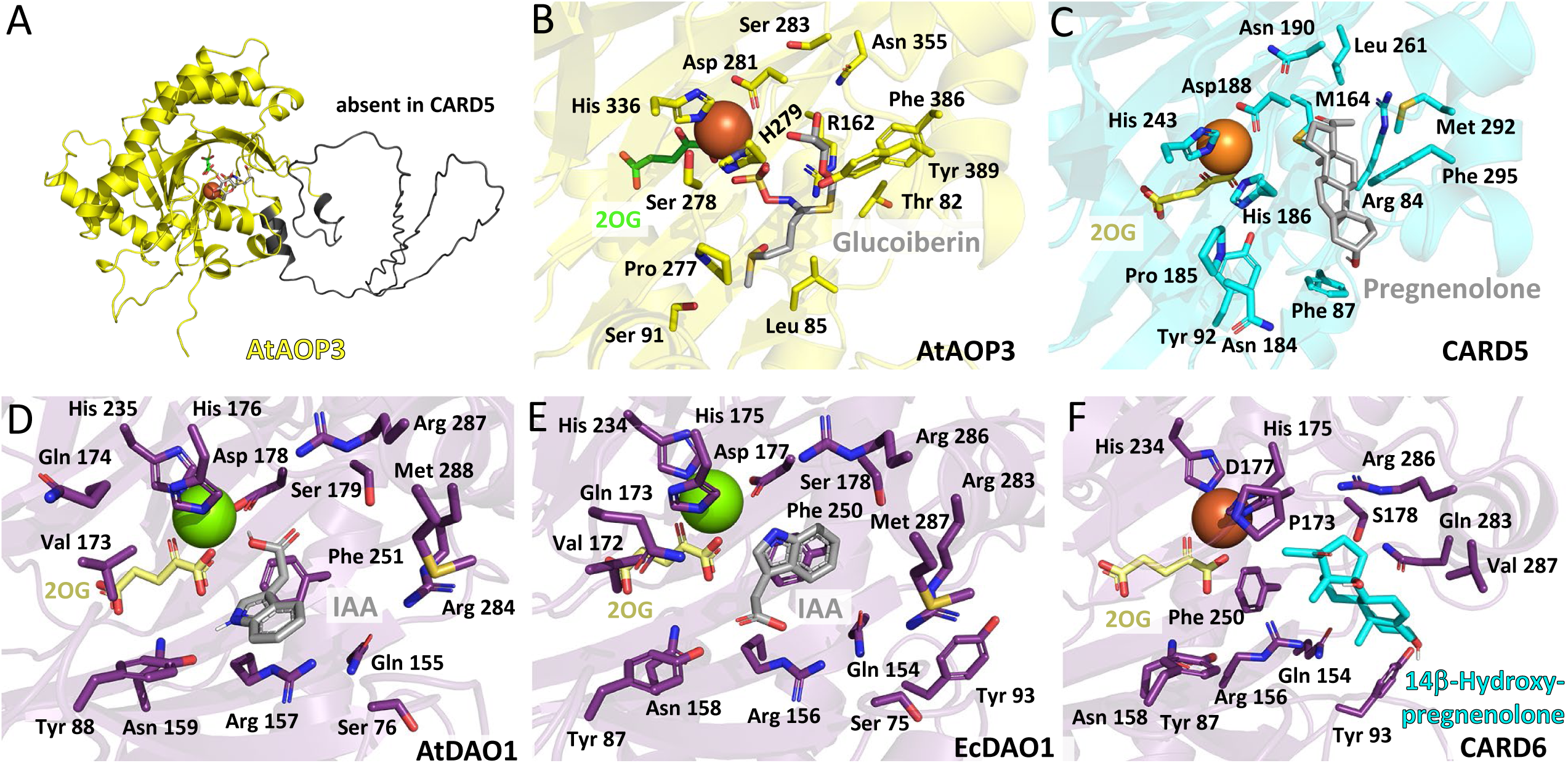
Substrate docking and analysis of residues involved in substrate binding. (A) The *Arabidopsis thaliana* AOP3 (yellow) contains an extra 95 amino acids (black) that are absent in *Erysimum cheiranthoides* CARD5. (B) The AtAOP3 active site with Fe(II) (orange) and 2-oxoglutarate (2OG; green) bound. Glucoiberin (grey) was docked into the active site, and residues that coordinate the metal binding site and that comprise the active site but are variable between AtAOP3 and CARD5 are shown as sticks. (C) Pregnenalone **2** (grey) was docked into the CARD5 active site, and sticks are side chains are shown for the active site amino acids that are variable with AtAOP3. (D and E) The *A. thaliana* and *E. cheiranthoides* DAO1 active sites with indole-3-acetic acid (IAA; grey), Mg (green), and 2OG (yellow) have identical active site residues. (F) The CARD6 active site with Fe(II) (orange), 2OG, and 14β-hydroxypregnenolone **12** (cyan) reveals the residues that are conserved and variable in comparison to EcDAO1.

The *A. thaliana* DAO1 active site residues are entirely conserved in the *E. cheiranthoides* DAO1 (Figure 5D, E), suggesting that they are critical for DAO1’s function as the primary mechanism for IAA oxidative catabolism in plants (Zhang *et al*., 2016; Jin *et al*., 2020). There are few notable differences in the active sites of EcDAO1 and CARD6, with three active site variations that stand out as being important in the evolution of steroid recognition. The polar, charged Arg 283 in EcDAO1 is instead a smaller polar Gln in the CARD6 active site, and the polar Gln 173 in EcDAO1 is a nonpolar Pro 173 in CARD6 (Figure 5 E, F). The nonpolar Met 287 in EcDAO1 corresponds to Val 287 in CARD6, which has a smaller nonpolar side chain.

Taken together, these active site differences likely permit the binding of larger steroid substrates. When docked into the CARD6 active site, 14β-hydroxypregnenolone **12** is positioned for 21-hydroxylation.

## Discussion

### Identification of probable steroid 14β- and 21-hydroxylases

This study identified two key enzymes in cardenolide biosynthesis in *E. cheiranthoides*, CARD5 and CARD6, the probable steroid 14β- and 21-hydroxylases, respectively. Using gene knockouts, we show that both enzymes are required for cardenolide biosynthesis in *E. cheiranthoides*, and we clarify the order of hydroxylation, with 14β-hydroxylation likely occurring before 21-hydroxylation. Using these enzymes, we were able to partially reconstitute cardenolide biosynthesis in *N. benthamiana*. The discovery of these enzymes brings us two steps closer to engineering of the full cardenolide pathway in a heterologous system and adds two new pregnane-modifying enzymes to the toolkit for production of other important steroidal compounds. CARD5 is of particular interest, as no other enzymes capable of steroid 14β-hydroxylation are known. While the exact activity of these enzymes requires further characterization, this opens the door to chemo-enzymatic synthesis of diverse steroidal metabolites that may be useful in human health and treatment of disease (Zhao *et al*., 2022).

Based on the cardenolide chemotype of *card2*, *card3*, and *card4* mutants, which accumulate dehydrocardenolides (Younkin *et al*., 2024b), it is unsurprising that CARD5 and CARD6 are active on multiple intermediates in cardenolide biosynthesis. Whether or not cardenolide biosynthesis follows this grid-like pathway (Figure 4) *in planta*, or whether this is an artefact of partial pathway reconstruction in a heterologous system, is not known. Furthermore, the unknown structure of the products of CARD6 in the transient expression experiments, compounds **15**-**20**, complicates pathway elucidation. While it is clear that the glycosylation of these compounds is catalyzed by endogenous *N. benthamiana* enzymes, the exact activity of CARD6 in these remains ambiguous. For example, if the proposed structures are correct, it is possible that CARD6 hydroxylates compounds **12**-**14** at carbon 21, and an aldehyde or steroid dehydrogenase from *N. benthamiana* further oxidizes these intermediates to compounds **15**-**20**. Alternatively, CARD6 may catalyze the formation of these 21-oxo-pregnanes, and cardenolide biosynthesis requires a specialized aldehyde dehydrogenase from *E. cheiranthoides* to form compound **8** (Figure 1A). Future *in vitro* assays with purified recombinant enzymes and structural elucidation of intermediates via NMR will clarify these possibilities.

### On the mechanism for stereochemical inversion at C14

The stereochemical inversion at carbon 14 that apparently accompanies hydroxylation, which is critical for the biological activity of cardenolides, has been a matter of some interest (Kreis et al., 1998; Zhao et al., 2022). The identification of the probable 14β-hydroxylase as a 2OGD facilitates speculation as to the catalytic mechanism that allows for deprotonation and subsequent hydroxylation to occur on opposite faces of the pregnane substrate. In the consensus reaction mechanism for 2OGDs, a Fe(IV)-oxo intermediate that abstracts a proton from the primary substrate, forming a substrate radical (Martinez & Hausinger, 2015). For epipregnanolone, if this radical is formed via abstraction of H_15β_, a 1,2-hydride shift of H_14ɑ_ would form a more stable tertiary radical intermediate. Hydroxylation could then proceed from the same face of the molecule from which the proton was abstracted, while also resulting in the requisite inversion of stereochemistry at carbon 14. While this mechanism is highly speculative, such rearrangements during hydroxylation by 2OGDs are not unprecedented. For example, in benzoxazinoid biosynthesis in maize, conversion of DIMBOA-Glc to TRIMBOA-Glc involves rearrangement of a methoxy group during hydroxylation by BX13, a 2OGD (Handrick et al., 2016). Further studies to confirm the regio- and stereo-specificity of CARD5 and to investigate this critical stereochemical inversion are warranted.

### Gene duplication and neofunctionalization in the evolution of cardenolide biosynthesis

Examination of the genomic context and sequence evolution of *CARD5* and *CARD6* provides concrete examples of the processes underlying the evolution of novel phytochemicals. In both cases, we observe gene duplication followed by shifts in substrate preference and activity, but these two new case studies provide insight into variation in this process. The case of *CARD6* appears straightforward. In the ancestor of both *E. cheiranthoides* and *A. thaliana*, *DAO* underwent a tandem duplication, resulting in *DAO1* and *DAO2*. While there has likely been subfunctionalization between *DAO1* and *DAO2*, both appear to primarily function in the oxidative inactivation of indole acetic acid (Zhang *et al*., 2016). In *Erysimum*, *DAO1* was duplicated a second time (Figure 2A, B), with one copy undergoing a radical shift in substrate preference to accept 14β-hydroxypregnanes, eventually becoming *CARD6*. CARD6’s new role in cardenolide biosynthesis is likely the result of three variable active site residues (Pro 173, Gln 283, and Val 287) that allow the enzyme to accommodate larger steroid substrates (Figure 5E, F).

The evolutionary history of *CARD5* appears more complex. The GS-AOP locus in *A. thaliana* has been the subject of substantial interest due to the relationship between structural variation at the locus and glucosinolate diversity (Kliebenstein *et al*., 2001; Chan *et al*., 2010). While AtAOP2 and AtAOP3 are involved in glucosinolate modification, the function of AtAOP1 remains unknown. The evolutionary history in *Erysimum* is further complicated by four additional duplications of *AOP1* scattered across three clusters separated by several million base pairs (Figure 2C,D). Examination of macrosynteny in this genomic region reveals evidence of substantial rearrangements and chromosomal fission (Züst *et al*., 2020). At least one of these copies, *CARD5*, took on a new role essential to cardenolide metabolism. The functions of the other copies remain unknown in *Erysimum*, but it is possible that others may be involved in further steps in cardenolide biosynthesis or modification. In fact, two of the others, *Erche03g038390* and *Erche02g016180*, are co-expressed with cardenolide biosynthesis genes (Younkin *et al*., 2024b). The close relationship of this clade to *Sl23DOX* (Figure 2C) hints that this group of 2OGDs may readily accept steroidal substrates, but a more thorough investigation of activity across DOXC20 2OGD family is needed before conclusions can be drawn. It is intriguing to find that closely related paralogous genes, *AOP2*/*AOP3* and *CARD5*, evolved key functions in the biosynthesis of structurally dissimilar defensive metabolites, glucosinolates and cardenolides respectively. Further investigation into the ancestral function of this clade and subsequent shifts in substrate preference will provide insights into how DOXC20 2OGDs provided the raw genetic material for the evolution of diverse defensive metabolites.

Furthermore, the ability of CARD5 to act on pregnenolone hints at a minimal pathway for plant metabolites with cardiotonic activity. By expressing only *CARD1-CARD5*, in *N. benthamiana*, we see the accumulation of compound **14**, which is likely 14β-hydroxyepipregnanolone. CARD5 is also active on pregnenolone **2**, so it is likely that compound **12** could be produced by the expression of only *CARD1* and *CARD5*. According to studies on the structure-activity relationship of steroidal compounds, these structures are sufficient for at least some inhibitory activity against Na^+^/K^+^-ATPases (Schönfeld *et al*., 1985). Production of low levels of pregnane metabolites such as pregnenolone and progesterone is widespread in plants (Iino *et al*., 2007; Lindemann, 2015), so an increase in pregnane production coupled with the evolution of 14β-hydroxylase activity may be sufficient to provide an ecological defense against insects. After production of 14β-hydroxypregnanes is established, further modifications to this minimal structure, including lactone ring formation and glycosylation would strengthen and fine-tune biological activity. This model of a step-wise pathway evolution, with only small, biochemically accessible changes required for production of an active metabolite may help explain the recurrent evolution of cardenolides as defensive compounds across land plants and some animals (Agrawal *et al*., 2012).

## Acknowledgements

This research was funded by the United States Department of Agriculture (2020-67013-30896 to G.J.), the US National Science Foundation (DGE-2139899 to G.C.Y., IOS-1645256 to G.J. and MCB-2214883 to C.K.H.), Williams College (C.K.H), the Triad Foundation (G.J.), and a Chemistry Biology Interface Training Program fellowship from the National Institutes of Health/National Institute of General Medical Sciences (T32GM138826).

## Author Contributions

GCY and GJ designed the research; GCyand CKH performed the research; GCY and CKH analyzed data; GCY, CKH, and GJ wrote and edited the manuscript.

## Data Availability

The raw data that support the findings of this study are available in the Supporting Information. Seeds from mutant lines are available from the Arabidopsis Biological Resource Center (ABRC; https://abrc.osu.edu/).

---

**Figure S1.**
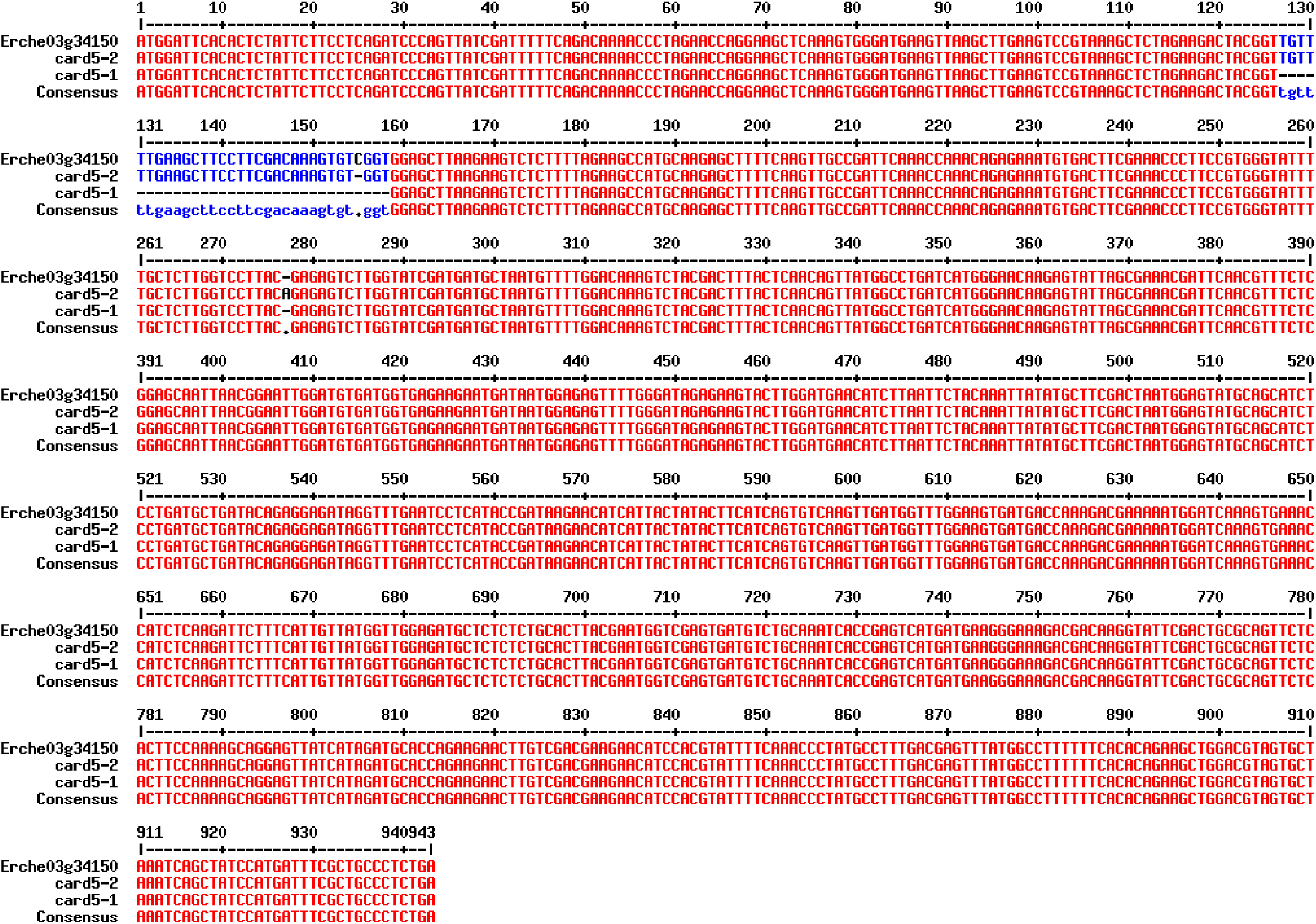
Aligned nucleotide coding sequences of *Erche03g034150* (*CARDENOLIDE METABOLISM 5*; *CARD5*) from wildtype *E. cheiranthoides* and mutant lines. *card5-1* and *card5-2* were generated with CRISPR/Cas9. Sequences of gRNAs used for generation of these lines are available in Table S1. MultAlin (http://multalin.toulouse.inra.fr/multalin/) was used to produce the alignment.

**Figure S2.**
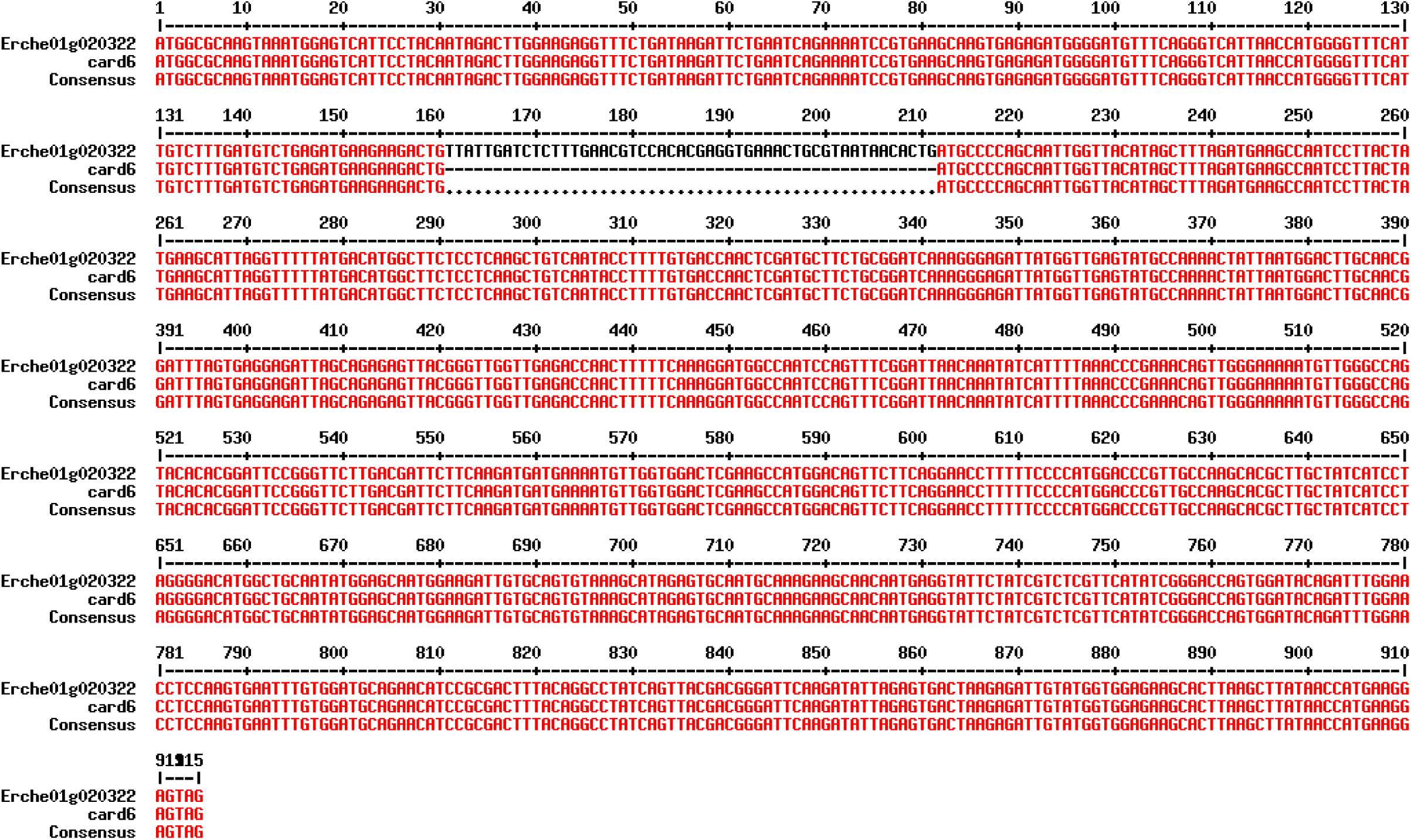
Aligned nucleotide coding sequences of *Erche01g020322* (*CARDENOLIDE METABOLISM 6*; *CARD6*) from wildtype *E. cheiranthoides* and a mutant line. *card6* was generated with CRISPR/Cas9. Sequences of gRNAs used for generation of these lines are available in Table S1. MultAlin (http://multalin.toulouse.inra.fr/multalin/) was used to produce the alignment.

**Figure S3.**
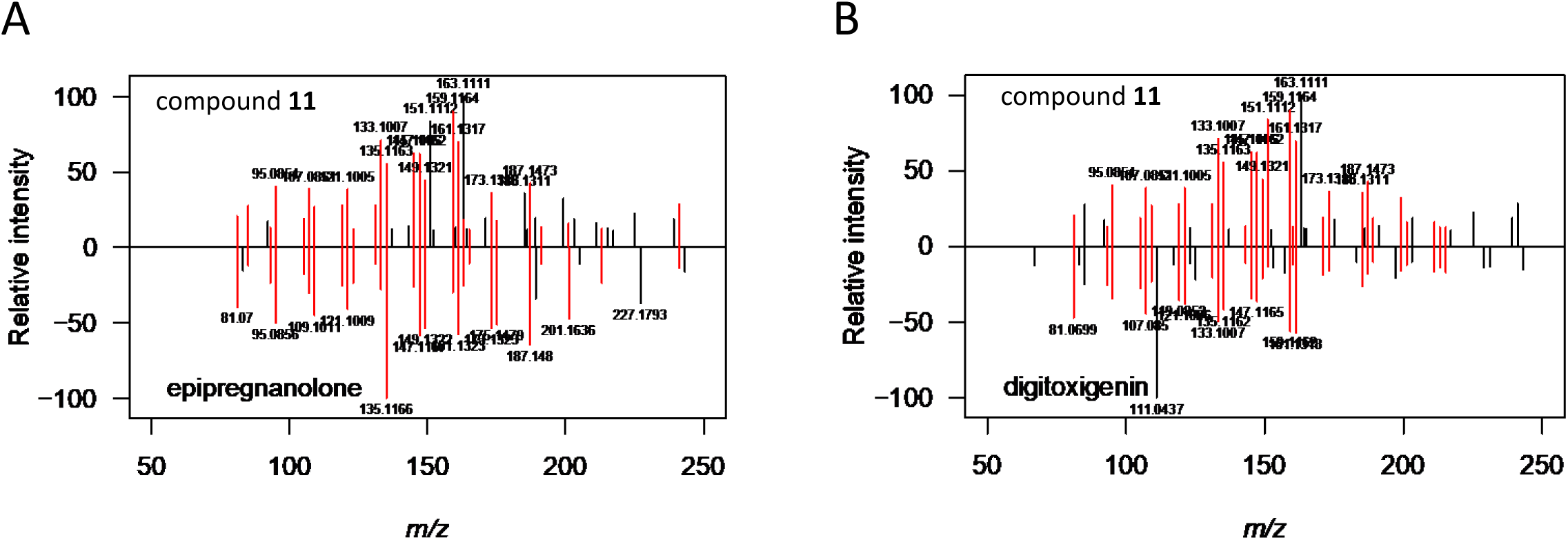
MSMS spectrum for compound 11 from *card6* mutant line. MSMS spectrum for compound **11**, a glycosylated cardenolide intermediate that accumulates to high levels in the *card6 Erysimum cheiranthoides* line compared with (A) epipregnanolone and (B) digitoxigenin standards. For compound **11**, [M+H]^+^ is absent, and [M+Na]^+^ does not fragment well during MSMS. In order to facilitate comparison to digitoxigenin and epipregnanolone spectra, an MS fragment corresponding to the neutral loss of the glycoside was selected for MSMS analysis: [M-deoxyhexose-H2O+H]^+^ (*m/z*=317.2481). Similarly, [M-H2O+H]^+^ was selected for both digitoxigenin (*m/z*=357.2430) and epipregnanolone (*m/z*=301.2531) standards. Peaks between *m/z*=50-250 are shown, with peaks shared between mirrored spectra within 0.005 Da colored in red.

**Figure S4.**
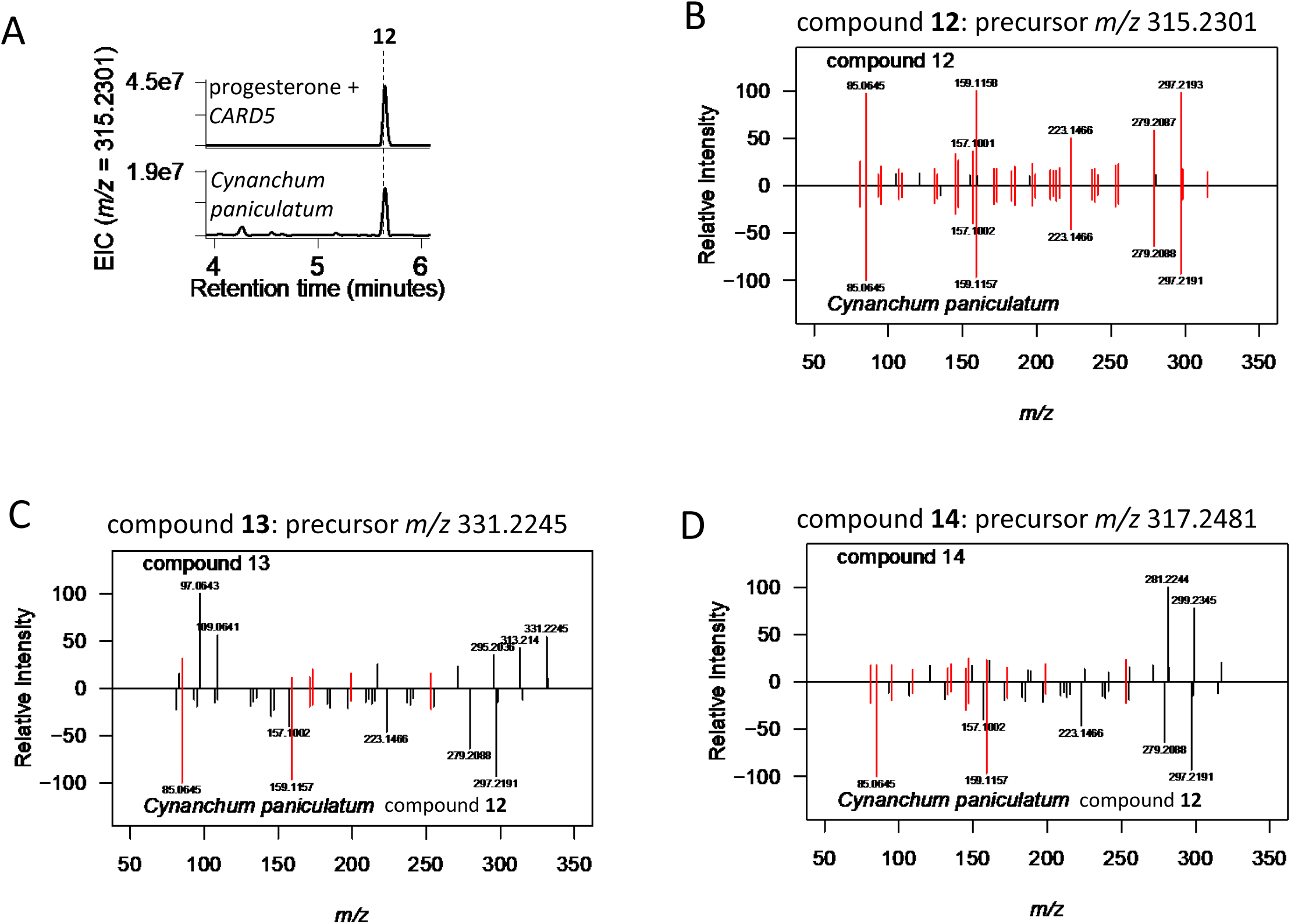
MSMS spectra for compounds 12-14 from substrate feeding upon co-infiltration with *CARD5* in *Nicotiana benthamiana*. These three compounds are hypothesized to be 14β-hydroxy derivatives of pregnenolone (compound **12**), progesterone (compound **13**), and epipregnanolone (compound **14**), differing only in the presence of a 3-oxo- or 3-hydroxy group, and saturation at carbon 5. 14β-hydroxypregnenolone (also called ketocalogenin) has been identified from *Cynanchum paniculatum* via purification and NMR. Consistent with the hypothesis that compound **12** is 14β-hydroxypregnenolone, compound **12** is also found in *C. paniculatum* extracts (A,B). Compounds **13** and **14** are predicted to differ from compound **12** by the presence or absence of a double bond. Therefore, some MSMS peaks are expected to be shifted by 2-4 Da between these spectra (C,D). For compound **12** from both *C. paniculatum* and *N. benthamiana* (B) and compound **14** (D), [M-H20+H]^+^ was selected for fragmentation because [M+H]^+^ was not detected, and [M+Na]^+^ did not fragment well. [M+H]^+^ was used as the precursor ion for compound **13** (B). Peaks shared between mirrored spectra within 0.005 Da colored in red.

**Figure S5.**
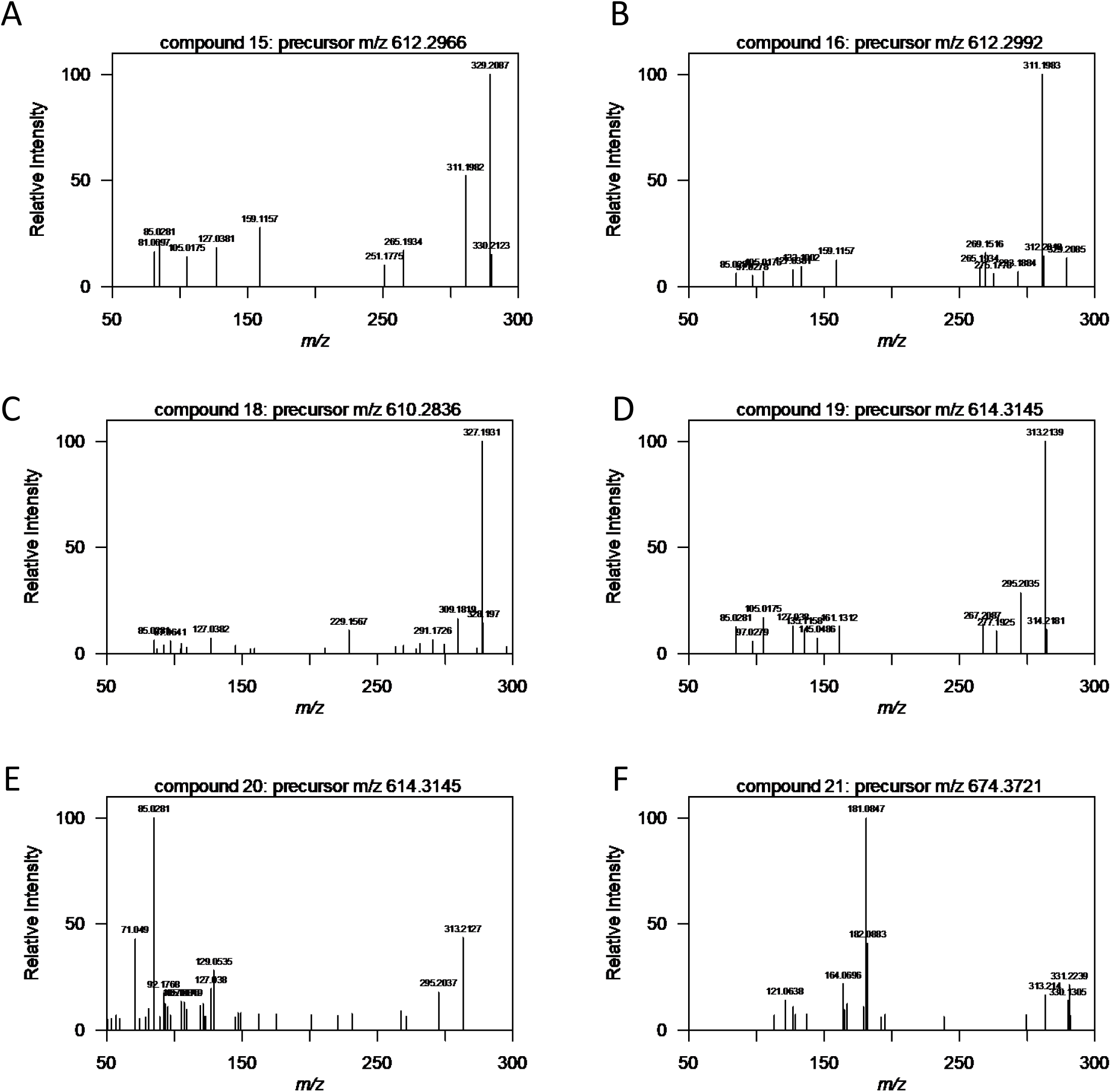
MSMS spectra for compounds 15-21 from substrate feeding of CARD5 + CARD6 in *Nicotiana benthamiana*. These three compounds are hypothesized to be 14β-hydroxy, 21-oxo derivatives of pregnenolone (compounds **15** and **16**), progesterone (compound **18**—no spectrum was collected for compound **17**), epipregnanolone (compounds **19** and **20**), differing only based on the presence of a 3-oxo- or 3-hydroxy group, and saturation at carbon 5. Compound **21** is the oxidation product of compound **11** by CARD6. Many peaks are expected to be shifted by 2-4 Da between these spectra, including the peaks 309/327 for compound **18**, 311/329 for compounds **15** and **16**, and 313/331 for compounds **19**, **20**, and **21**. For compound **21** (F), the strong peak at 181 Da corresponds to a hexose. [M-NH4]^+^ was used as the precursor ion for fragmentation of all compounds because [M+H]^+^ was not detected, and [M+Na]^+^ did not fragment well.

## REFERENCES

Agrawal AA, Petschenka G, Bingham RA, Weber MG, Rasmann S. 2012. Toxic cardenolides: Chemical ecology and coevolution of specialized plant-herbivore interactions. New Phytologist 194: 28–45.

Alani ML, Younkin GC, Mirzaei M, Kumar P, Jander G. 2021. Acropetal and basipetal cardenolide transport in *Erysimum cheiranthoides* (wormseed wallflower). Phytochemistry 192: 112965.

Amor M, Parker KL, Globerman H, New MI, White PC. 1988. Mutation in the *CYP21B* gene (Ile-172→Asn) causes steroid 21-hydroxylase deficiency. Proceedings of the National Academy of Sciences of the United States of America 85: 1600–1604.

Benton HP, Want EJ, Ebbels TMD. 2010. Correction of mass calibration gaps in liquid chromatography-mass spectrometry metabolomics data. Bioinformatics 26: 2488–2489.

Berardini TZ, Reiser L, Li D, Mezheritsky Y, Muller R, Strait E, Huala E. 2015. The Arabidopsis information resource: Making and mining the ‘gold standard’ annotated reference plant genome. Genesis 53: 474–485.

Bose D, Elliott D, Kobayashi T, Templeton JF, Kumar VPS, LaBella FS. 1988. 14β-Hydroxyprogesterone binds to the digitalis receptor, inhibits the sodium pump and enhances cardiac contractility. British Journal of Pharmacology 93: 453–461.

Carroll E, Gopal BR, Raghavan I, Mukherjee M, Wang ZQ. 2023. A cytochrome P450 CYP87A4 imparts sterol side-chain cleavage in digoxin biosynthesis. Nature Communications 14: 4042.

Chan EKF, Rowe HC, Kliebenstein DJ. 2010. Understanding the evolution of defense metabolites in *Arabidopsis thaliana* using genome-wide association mapping. Genetics 185: 991–1007.

Chen J, Tang J, Xi Y, Dai Z, Bi C, Chen X, Fan F, Zhang X. 2019. Production of 14α-hydroxysteroids by a recombinant *Saccharomyces cerevisiae* biocatalyst expressing of a fungal steroid 14α-hydroxylation system. Applied Microbiology and Biotechnology 103: 8363–8374.

Deluca ME, Seldes AM, Gros EG. 1987. The 14ß-HydroxyIation in the Biosynthesis of Cardenolides in Digitalis purpurea. The Role of 3ß-Hydroxy-5ß-pregn-8(14)-en-20-one. Zeitschrift für Naturforschung 42c: 77–78.

Forli S, Huey R, Pique ME, Sanner MF, Goodsell DS, Olson AJ. 2016. Computational protein-ligand docking and virtual drug screening with the AutoDock suite. Nature Protocols 11: 905–919.

Gatto L, Gibb S, Rainer J. 2021. MSnbase, Efficient and Elegant R-Based Processing and Visualization of Raw Mass Spectrometry Data. Journal of Proteome Research 20: 1063–1069.

Gatto L, Lilley K. 2012. MSnbase - an R/Bioconductor package for isobaric tagged mass spectrometry data visualization, processing and quantitation. Bioinformatics 28: 288–289.

Graves S, Piepho H-P, Dorai-Raj S, Selzer L. 2023. multcompView: Visualizations of Paired Comparisons.

Hagel JM, Facchini PJ. 2018. Expanding the roles for 2-oxoglutarate-dependent oxygenases in plant metabolism. Natural Product Reports 35: 721–734.

Handrick V, Robert CAM, Ahern KR, Zhou S, Machado RAR, Maag D, Glauser G, Fernandez-Penny FE, Chandran JN, Rodgers-Melnik E, et al. 2016. Biosynthesis of 8-O-methylated benzoxazinoid defense compounds in maize. Plant Cell 28: 1682–1700.

Hekkelman ML, de Vries I, Joosten RP, Perrakis A. 2023. AlphaFill: enriching AlphaFold models with ligands and cofactors. Nature Methods 20: 205–213.

Hoang DT, Chernomor O, Von Haeseler A, Minh BQ, Vinh LS. 2018. UFBoot2: Improving the ultrafast bootstrap approximation. Molecular Biology and Evolution 35: 518–522.

Iino M, Nomura T, Tamaki Y, Yamada Y, Yoneyama K, Takeuchi Y, Mori M, Asami T, Nakano T, Yokota T. 2007. Progesterone: Its occurrence in plants and involvement in plant growth. Phytochemistry 68: 1664–1673.

Jin S-H, Lee H, Shin Y, Kim J-H, Rhee S. 2020. Crystal structure of the indole-3-acetic acid-catabolizing enzyme DAO1 from *Arabidopsis thaliana*. Journal of Structural Biology 212: 107632.

Jumper J, Evans R, Pritzel A, Green T, Figurnov M, Ronneberger O, Tunyasuvunakool K, Bates R, Žídek A, Potapenko A, et al. 2021. Highly accurate protein structure prediction with AlphaFold. Nature 596: 583–589.

Kawahara Y, de la Bastide M, Hamilton JP, Kanamori H, Mccombie WR, Ouyang S, Schwartz DC, Tanaka T, Wu J, Zhou S, et al. 2013. Improvement of the Oryza sativa nipponbare reference genome using next generation sequence and optical map data. Rice 6.

Kawai Y, Ono E, Mizutani M. 2014. Evolution and diversity of the 2-oxoglutarate-dependent dioxygenase superfamily in plants. Plant Journal 78: 328–343.

Klein J. 2024. Progesterone Metabolism in Digitalis and Other Plants-60 Years of Research and Recent Results. Plant & Cell Physiology 65: 1500–1514.

Klein J, Horn E, Ernst M, Leykauf T, Leupold T, Dorfner M, Wolf L, Ignatova A, Kreis W, Munkert J. 2021. RNAi-mediated gene knockdown of progesterone 5β-reductases in *Digitalis lanata* reduces 5β-cardenolide content. Plant Cell Reports 40: 1631–1646.

Kliebenstein DJ, Lambrix VM, Reichelt M, Gershenzon J, Mitchell-Olds T. 2001. Gene duplication in the diversification of secondary metabolism: Tandem 2-oxoglutarate-dependent dioxygenases control glucosinolate biosynthesis in Arabidopsis. Plant Cell 13: 681–693.

Kreis W. 2017. The Foxgloves (*Digitalis*) Revisited. Planta Medica 83: 962–976.

Kreis W, Hensel A, Stuhlemmer U. 1998. Cardenolide Biosynthesis in Foxglove. Planta Medica 64: 491–499.

Kunert M, Langley C, Lucier R, Ploss K, Rodríguez López CE, Serna Guerrero DA, Rothe E, O’Connor SE, Sonawane PD. 2023. Promiscuous CYP87A enzyme activity initiates cardenolide biosynthesis in plants. Nature Plants 9: 1607–1617.

Leykauf T, Klein J, Ernst M, Dorfner M, Ignatova A, Kreis W, Lanig H, Munkert J. 2023. Overexpression and RNAi-mediated knockdown of two 3β-hydroxy-Δ5-steroid dehydrogenase genes in Digitalis lanata shoot cultures reveal their role in cardenolide biosynthesis. Planta Medica 89: 833–847.

Lindemann P. 2015. Steroidogenesis in plants - Biosynthesis and conversions of progesterone and other pregnane derivatives. Steroids 103: 145–152.

Madeira F, Pearce M, Tivey ARN, Basutkar P, Lee J, Edbali O, Madhusoodanan N, Kolesnikov A, Lopez R. 2022. Search and sequence analysis tools services from EMBL-EBI in 2022. Nucleic Acids Research 50: W276–W279.

Martinez S, Hausinger RP. 2015. Catalytic mechanisms of Fe(II)- and 2-Oxoglutarate-dependent oxygenases. Journal of Biological Chemistry 290: 20702–20711.

Minh BQ, Schmidt HA, Chernomor O, Schrempf D, Woodhams MD, Von Haeseler A, Lanfear R, Teeling E. 2020. IQ-TREE 2: New Models and Efficient Methods for Phylogenetic Inference in the Genomic Era. Molecular Biology and Evolution 37: 1530–1534.

Mirdita M, Schütze K, Moriwaki Y, Heo L, Ovchinnikov S, Steinegger M. 2022. ColabFold: making protein folding accessible to all. Nature Methods 19: 679–682.

Mirzaei M, Younkin GC, Powell AF, Alani ML, Strickler SR, Jander G. 2024. Aphid Resistance Segregates Independently of Cardenolide and Glucosinolate Content in an *Erysimum cheiranthoides* (Wormseed Wallflower) F2 Population. Plants 13: 466.

Nakayasu M, Akiyama R, Kobayashi M, Lee HJ, Kawasaki T, Watanabe B, Urakawa S, Kato J, Sugimoto Y, Iijima Y, et al. 2020. Identification of α-tomatine 23-hydroxylase involved in the detoxification of a bitter glycoalkaloid. Plant and Cell Physiology 61: 21–28.

Nakayasu M, Umemoto N, Ohyama K, Fujimoto Y, Lee HJ, Watanabe B, Muranaka T, Saito K, Sugimoto Y, Mizutani M. 2017. A dioxygenase catalyzes steroid 16α-hydroxylation in steroidal glycoalkaloid biosynthesis. Plant Physiology 175: 120–133.

Nelson D, Werck-Reichhart D. 2011. A P450-centric view of plant evolution. Plant Journal 66: 194–211.

Newman RA, Yang P, Pawlus AD, Block KI. 2008. Cardiac glycosides as novel cancer therapeutic agents. Molecular Interventions 8: 36–49.

Norn S, Kruse PR. 2004. Cardiac glycosides: From ancient history through Withering’s foxglove to endogenous cardiac glycosides. Dansk Medicinhistorisk Arbog: 119–132.

Ohnishi T, Yokota T, Mizutani M. 2009. Insights into the function and evolution of P450s in plant steroid metabolism. Phytochemistry 70: 1918–1929.

Porco S, Pěnčík A, Rasheda A, Vo U, Casanova-Sáez R, Bishopp A, Golebiowska A, Bhosale R, Swarupa R, Swarup K, et al. 2016. Dioxygenase-encoding AtDAO1 gene controls IAA oxidation and homeostasis in *Arabidopsis*. Proceedings of the National Academy of Sciences of the United States of America 113: 11016–11021.

R Core Team. 2020. R: A Language and Environment for Statistical Computing.

Renwick J, Radke C, K S-G. 1989. Chemical consituents of *Erysimum cheiranthoides* deterring oviposition by cabbage butterfly, *Pieris rapae*. Journal of Chemical Ecology 15: 2161–2169.

Schönfeld W, Weiland J, Lindig C, Masnyk M, Kabat MM, Kurek A, Wicha J, Repke KRH. 1985. The lead structure in cardiac glycosides is 5β,14β-androstane-3β,14-diol. Naunyn-Schmiedeberg’s Archives of Pharmacology 329: 414–426.

Sievers F, Wilm A, Dineen D, Gibson TJ, Karplus K, Li W, Lopez R, McWilliam H, Remmert M, Söding J, et al. 2011. Fast, scalable generation of high-quality protein multiple sequence alignments using Clustal Omega. Molecular Systems Biology 7.

Smith CA, Want EJ, O’Maille G, Abagyan R, Siuzdak G. 2006. XCMS: Processing mass spectrometry data for metabolite profiling using nonlinear peak alignment, matching, and identification. Analytical Chemistry 78: 779–787.

Sugama K, Hayashi K, Mitsuhashi H, Kaneko K. 1986. Studies on the contituents of Asclepiadaceae Plants. LXVI. The structrues of three new glycosides, cynaponosides A, B, and C, from the Chinese drug ‘Xu-Chang-Qing,’ Cynanchum paniculatum KITAGAWA. Chemical Pharmaceutical Bulletin 34: 4500–4507.

Suzuki K, Sanga K, Chikaoka Y, Itagaki E. 1993. Purification and properties of cytochrome P-450 (P-450lun) catalyzing steroid 11β-hydroxylation in *Curvularia lunata*. Biochimica et Biophysica Acta 1203: 215–223.

Tautenhahn R, Bottcher C, Neumann S. 2008. Highly sensitive feature detection for high resolution LC/MS. BMC Bioinformatics 9: 504.

Tello-Ruiz MK, Jaiswal P, Ware D. 2022. Gramene: A Resource for Comparative Analysis of Plants Genomes and Pathways. In: Edwards D, ed. Plant Bioinformatics. New York, NY: Springer US, 101–131.

o 2016. W-IQ-TREE: a fast online phylogenetic tool for maximum likelihood analysis. Nucleic Acids Research 44: W232–W235.

Trott O, Olson AJ. 2010. AutoDock Vina: Improving the speed and accuracy of docking with a new scoring function, efficient optimization, and multithreading. Journal of Computational Chemistry 31: 455–461.

White PC, New MI, Dupont B. 1986. Structure of human steroid 21-hydroxylase genes. Proceedings of the National Academy of Sciences of the United States of America 83: 5111– 5115.

Wong RW, Lingwood CA, Ostrowski MA, Cabral T, Cochrane A. 2018. Cardiac glycoside/aglycones inhibit HIV-1 gene expression by a mechanism requiring MEK1/2-ERK1/2 signaling. Scientific Reports 8: 1–17.

Yasumoto S, Fukushima EO, Seki H, Muranaka T. 2016. Novel triterpene oxidizing activity of *Arabidopsis thaliana* CYP716A subfamily enzymes. FEBS Letters 590: 533–540.

Younkin GC, Alani ML, Páez-Capador A, Fischer HD, Mirzaei M, Hastings AP, Agrawal AA, Jander G. 2024a. Cardiac glycosides protect wormseed wallflower (*Erysimum cheiranthoides*) against some, but not all, glucosinolate-adapted herbivores. New Phytologist 242: 2719–2733.

Younkin G, Alani ML, Züst T, Jander G. 2024b. Four enzymes control natural variation in the steroid core of *Erysimum* cardenolides. bioRxiv.

Zalucki MP, Malcolm SB, Pauste TD, Hanlon CC, Brower LP, Clarke AR. 2001. It’s the first bites that count: Survival of first-instar monarchs on milkweeds. Austral Ecology 26: 547–555.

Zhang J, Lin JE, Harris C, Mastrotti Pereira FC, Wu F, Blakeslee JJ, Peer WA. 2016. DAO1 catalyzes temporal and tissue-specific oxidative inactivation of auxin in *Arabidopsis thaliana*. Proceedings of the National Academy of Sciences of the United States of America 113: 11010–11015.

Zhao Z, Zhang Y, Liu X, Zhang X, Liu S, Yu X, Ren Y, Zheng X, Zhou K, Jiang L, et al. 2013. A Role for a Dioxygenase in Auxin Metabolism and Reproductive Development in Rice. Developmental Cell 27: 113–122.

Zhao Y, Zhang B, Sun ZQ, Zhang H, Wang W, Wang ZR, Guo ZK, Yu S, Tan RX, Ge HM. 2022. Biocatalytic C14-Hydroxylation on Androstenedione Enabled Modular Synthesis of Cardiotonic Steroids. ACS Catalysis 12: 9839–9845.

Zhou Y, Ma Y, Zeng J, Duan L, Xue X, Wang H, Lin T, Liu Z, Zeng K, Zhong Y, et al. 2016. Convergence and divergence of bitterness biosynthesis and regulation in Cucurbitaceae. Nature Plants 2: 1–8.

Züst T, Strickler SR, Powell AF, Mabry ME, An H, Mirzaei M, York T, Holland CK, Kumar P, Erb M, et al. 2020. Independent evolution of ancestral and novel defenses in a genus of toxic plants (*Erysimum*, Brassicaceae). eLife 9: 1–42.

